# Oncostatin M is a Master Regulator of an Inflammatory Network in *Dnmt3a*-Mutant Hematopoietic Stem Cells

**DOI:** 10.1101/2023.07.12.548764

**Authors:** Logan S. Schwartz, Kira A. Young, Timothy M. Stearns, Nathan Boyer, Kristina D. Mujica, Jennifer J. Trowbridge

## Abstract

Age-associated clonal hematopoiesis (CH) occurs due to somatic mutations accrued in hematopoietic stem cells (HSCs) that confer a selective advantage in the context of aging. The mechanisms by which CH-mutant HSCs gain this advantage with aging are not comprehensively understood. Using unbiased transcriptomic approaches, we identify Oncostatin M (OSM) signaling as a candidate contributor to aging-driven *Dnmt3a*-mutant CH. We find that *Dnmt3a*-mutant HSCs from young mice do not functionally respond to acute OSM stimulation with respect to proliferation, apoptosis, hematopoietic engraftment, or myeloid differentiation. However, young *Dnmt3a*-mutant HSCs transcriptionally upregulate an inflammatory cytokine network in response to acute OSM stimulation including genes encoding IL-6, IL-1β and TNFα. In addition, OSM-stimulated *Dnmt3a*-mutant HSCs upregulate the anti-inflammatory genes *Socs3, Atf3* and *Nr4a1*, creating a negative feedback loop limiting sustained activation of the inflammatory network. In the context of an aged bone marrow (BM) microenvironment with chronically elevated levels of OSM, *Dnmt3a*-mutant HSCs upregulate pro-inflammatory genes but do not upregulate *Socs3, Atf3* and *Nr4a1*. Together, our work suggests that chronic inflammation with aging exhausts the regulatory mechanisms in young CH-mutant HSCs that resolve inflammatory states, and that OSM is a master regulator of an inflammatory network that contributes to age-associated CH.

## INTRODUCTION

The process of aging has a profound impact on tissues and cells throughout the body. In the hematopoietic stem and progenitor cell (HSPC) compartment, aging is accompanied by acquisition of somatic mutations. Given the long-lived nature of the HSPC pool, these mutations can be propagated for many cellular generations and through most mature hematopoietic cell progeny. While most of these somatic mutations are considered ‘neutral’, a subset can confer a selective growth advantage to HSPCs, leading to a condition termed age-associated clonal hematopoiesis (CH). The most common CH mutations are found in a subset of genes that encode canonical epigenetic or chromatin regulatory proteins such as DNA methyltransferase 3A (DNMT3A), tet methylcytosine dioxygenase 2 (TET2) and additional sex combs like-1 (ASXL1)^1, 2^. While CH is not a disease, it is associated with increased risk of age-associated pathologies such as blood cancers and ischemic stroke^1–7^. Understanding the cellular and molecular mechanisms underlying the selective advantage of HSPCs carrying these somatic mutations, and how and in whom this is a risk factor for age-associated disease, will further inform our understanding of CH-associated conditions.

By creating and utilizing genetically engineered mouse models, many research groups have reported functional and molecular consequences of the most common CH mutations in *DNMT3A*, *TET2*, and *ASXL1*^1, 2, 7–9^. In some of these studies, chronic stress and inflammation were found to act as selective pressures contributing to expansion of CH and development of CH-associated pathologies^1, 10, 11^. CH-mutant HSCs have different molecular and functional responses to pro-inflammatory cytokine signaling^12, 13^, and recent work suggests that CH-mutant hematopoietic cells can induce and maintain a pro-inflammatory state^14^. For example, *Dnmt3a*-mutant hematopoiesis is associated with elevated levels of IFNγ^15, 16^, IL-6^17^ and TNFα^18^. *Tet2*-mutant hematopoiesis is associated with elevated levels of IL-6^19^, IL-8^20^, IL-1β^21–23^ and TNFα^24^. *Asxl1*-mutant CH is associated with increased IFNγ and TNFα^25^. The extent to which multiple cytokines activate the same underlying inflammatory network contributing to CH and CH-associated pathologies, or whether individual cytokines have context-dependent effects, remains unknown. Addressing this gap in knowledge is important when considering therapeutic interventions that will be most effective in individuals at risk of CH-associated pathologies.

Using a Cre-inducible mouse model that our laboratory previously engineered to express a specific *Dnmt3a* mutation associated with CH and acute myeloid leukemia (Dnmt3a^R878H^)^26^, we found that placing these cells in a middle-aged BM microenvironment in the context of elevated pro-inflammatory cytokines promoted their selective advantage^18^. We discovered that TNFα:TNFR1 signaling was a key mediator of the selective advantage of *Dnmt3a-*mutant HSCs in the context of a middle-aged BM microenvironment^18^. However, as described above, TNFα is not the only cytokine known to contribute to *Dnmt3a*-mutant CH. Here, we further explored our transcriptomics data to identify and functionally test the role of other pro-inflammatory cytokines not yet evaluated in the context of *Dnmt3a*-mutant CH.

## RESULTS

### *Dnmt3a*-Mutant HSCs Activate Oncostatin M (OSM) Signaling in an Aged Bone Marrow Microenvironment

Our previously published work profiled molecular signatures associated *Dnmt3a*-mutant (R878H/+) hematopoiesis in a middle-aged BM microenvironment by performing RNA sequencing (RNA-seq) on independent biological replicates of control and *Dnmt3a*-mutant HSCs re-isolated from young and middle-aged recipient mice^18^ (Figure 1A). In new analysis of these data, we observed enrichment of a hallmark inflammatory response signature in *Dnmt3a*-mutant HSCs compared to control HSCs in middle-aged recipient mice but not in young recipient mice (Figure 1B), suggesting that a middle-aged environment promotes transcriptional responses in *Dnmt3a*-mutant HSCs to inflammatory factors. Examining the top leading-edge genes of this signature enrichment, we observed increased expression of the IFNγ-regulated genes *Bst2, Klf6* and *Icam1,* the inflammatory-responsive receptors *Tlr2* and *Ccrl2*, and the copper transporter *Slc31a2*. *Osm* was the only leading-edge gene encoding a secreted molecule, Oncostatin M (OSM), which is an IL-6 family cytokine known to be involved in the immunopathogenesis of solid tumors and myeloma ^27–31^. Given this observation, we hypothesized that increased expression of *Osm* results in a feed-forward loop of activation of the OSM signaling pathway.

**Figure 1.**
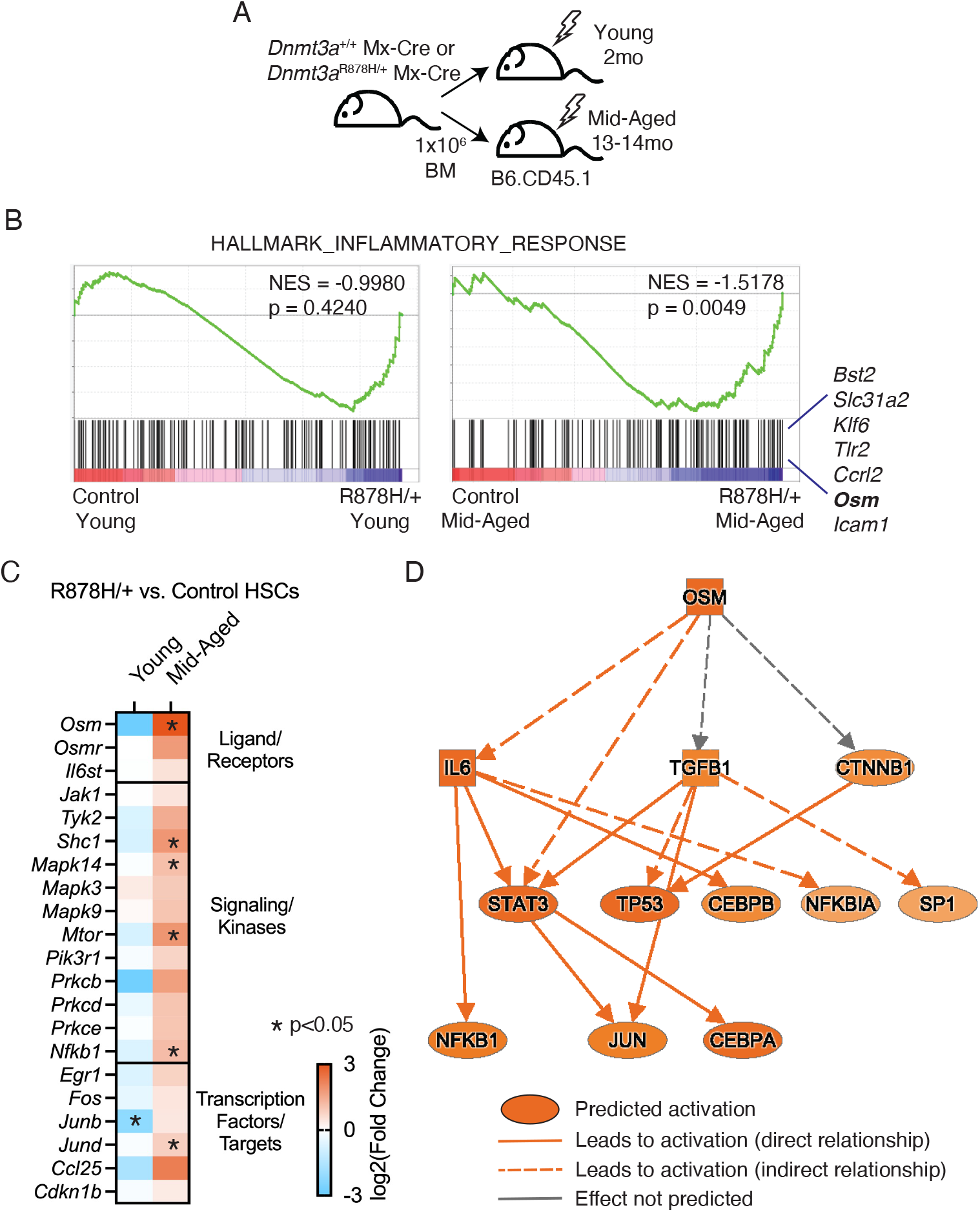
*Dnmt3a-*mutant HSCs upregulate *Osm* and OSM signaling genes in a middle-aged BM environment. (**A**) Schematic of transplant design into young (2mo) and middle-aged (13-14mo) recipient mice. HSCs (Lin^-^ c-Kit+ Sca-1^+^ Flt3^-^ CD150^+^ CD48^-^) were isolated at 4 months post-transplant and used for RNA-sequencing^18^. *n* = 3-4 biological replicates. (**B**) Gene set enrichment analysis of a hallmark inflammatory response signature in control vs. *Dnmt3a*-mutant (R878H/+) HSCs in young recipient mice (left) and in middle-aged recipient mice (right). (**C**) Heatmap of log2(FC) expression in OSM signaling pathway genes in *Dnmt3a*-mutant (R878H/+) compared to control HSCs in young (left column) and middle-aged (right column) recipient mice. \**P*<0.05. (**D**) Ingenuity pathway analysis showing predicted activation of OSM signaling in *Dnmt3a*-mutant (R878H/+) vs. control HSCs in middle-aged recipient mice.

To test this hypothesis, we interrogated the expression of a subset of genes known to be involved in and/or regulated by OSM signaling including transcripts encoding OSM signaling receptors (*Osmr, Il6st*), downstream kinases and signaling molecules (ex. *Shc1, Mapk14, Prkcb, Nfkb1*), and transcription factor targets (ex. *Egr1, Fos, Junb, Jund*). We observed trends toward increased expression and significant increase in expression (*P<*0.05) in most genes examined in *Dnmt3a*-mutant HSCs compared to control HSCs in middle-aged recipient mice but not in young recipient mice (Figure 1C). We used ingenuity pathway analysis (IPA) as a complementary approach to examine activation of OSM signaling in *Dnmt3a*-mutant HSCs compared to control HSCs in middle-aged recipient mice. We observed activation of an OSM-driven IL-6:STAT3 module in *Dnmt3a*-mutant HSCs in middle-aged recipient mice (Figure 1D). These results are consistent with activation of OSM signaling in *Dnmt3a*-mutant HSCs in the specific context of a middle-aged environment.

### Acute OSM Stimulation Does Not Impact Cell Cycle, Apoptosis, Proliferation or Myeloid Differentiation of Young *Dnmt3a*-Mutant HSPCs

Given that activation of OSM signaling is associated with expanded *Dnmt3a*-mutant hematopoiesis in an aged BM microenvironment, we hypothesized that OSM as a single stimulus would be sufficient to promote the selective advantage of young *Dnmt3a*-mutant HSPCs. Thus, we evaluated the effect of recombinant OSM on young *Dnmt3a*-mutant HSPC cycling, apoptosis, proliferation, and myeloid differentiation. We evaluated cell cycle status using Ki-67 and DAPI staining. Control and *Dnmt3a*-mutant HSPCs were prospectively isolated from young adult mice (3-6 months old) and stimulated overnight with 0, 100 or 500ng/ml recombinant murine OSM. Following flow cytometry analysis, cells were gated into G0 (Ki-67-DAPI-), G1 (Ki-67+ DAPI-) and S/G2/M (Ki-67+ DAPI+) fractions. We observed no significant differences in these proportions across any of the conditions, although there was a trend towards increased S/G2/M in *Dnmt3a*-mutant HSPCs with increasing doses of OSM (Figure 2A). We next evaluated apoptosis using Annexin V and PI staining. Control and *Dnmt3a*-mutant HSPCs were prepared and stimulated overnight as detailed above. Following flow cytometry analysis, cells were gated into live (Annexin V-PI-), early apoptotic (Annexin V+ PI-), late apoptotic (Annexin V+ PI+), and necrotic (Annexin V-PI+) fractions. We observed no significant differences in these proportions across any of the conditions (Figure 2B). Together, these data suggest that young control and *Dnmt3a*-mutant HSPCs do not respond to acute OSM stimulation with respect to altered cell cycle and apoptosis parameters.

**Figure 2.**
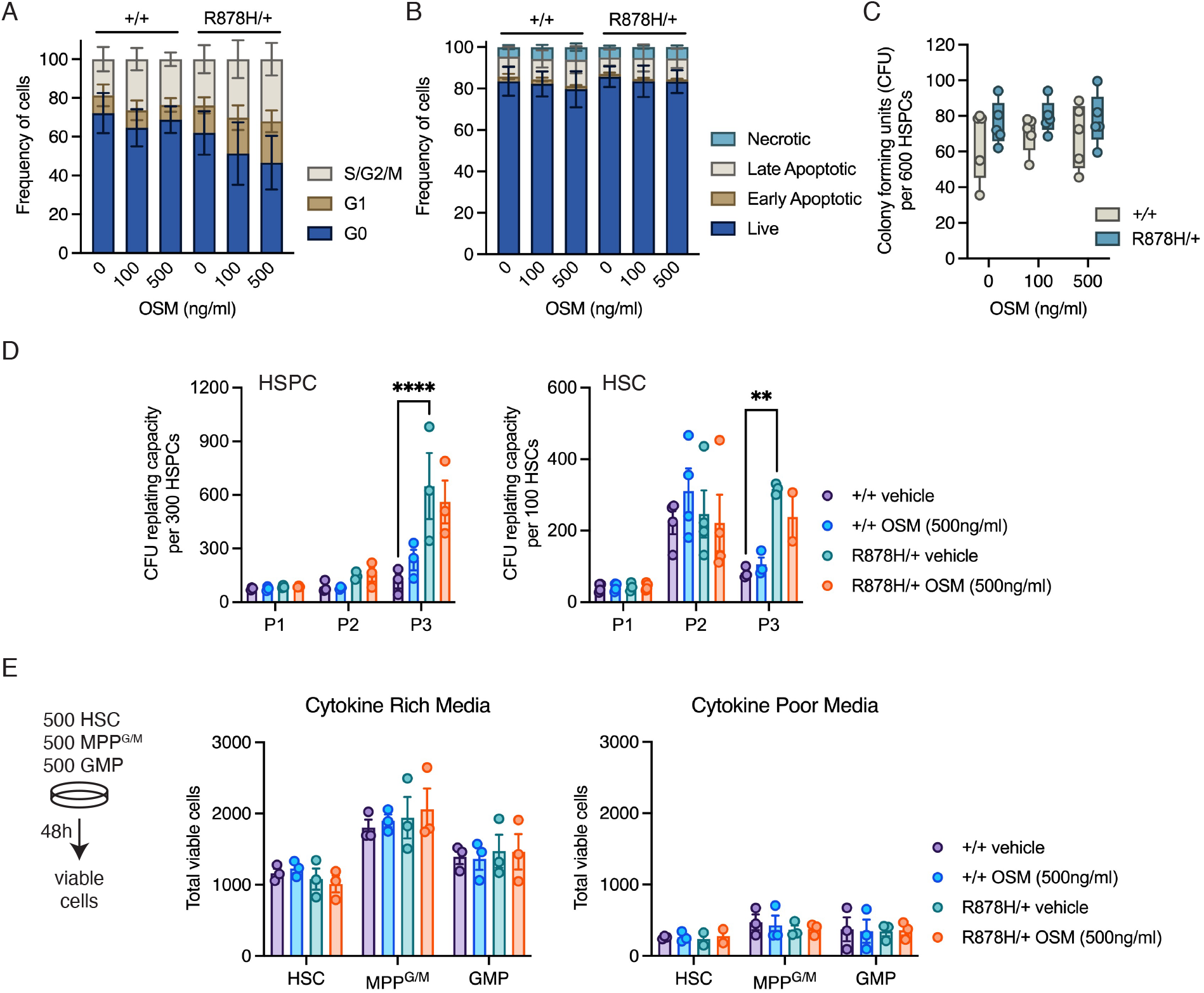
Cell cycle, apoptosis, proliferation, and myeloid differentiation in young *Dnmt3a*-mutant HSPCs are unaltered by acute OSM stimulation. **(A)** Frequency of control (+/+) and *Dnmt3a*-mutant (R878H/+) HSPCs in G0, G1, and S/G2/M cell cycle phases after stimulation with 0, 100, or 500ng/ml OSM for 24hrs. Bars represent mean ± SEM of *n* = 3. (**B**) Frequency of control (+/+) and *Dnmt3a*-mutant (R878H/+) HSPCs in live, early apoptotic, late apoptotic or necrotic gates after stimulation with 0, 100, or 500ng/ml OSM for 24hrs. Bars represent mean ± SEM of *n* = 3. (**C**) Colony-forming units (CFU) in control (+/+) and *Dnmt3a*-mutant (R878H/+) HSPCs with 0, 100 or 500ng/ml OSM. Dots show individual mice. *n* = 5. (**D**) Serial CFU in control (+/+) and *Dnmt3a*-mutant (R878H/+) HSPCs (left) and HSCs (right) with 0 or 500ng/ml OSM. Dots show individual mice, bars represent mean ± SEM of *n* = 3-4. \*\**p* < 0.01; \*\*\*\**p* < 0.0001 by two-way ANOVA with Tukey’s multiple comparisons test. (**E**) Total viable cells from control (+/+) and *Dnmt3a*-mutant (R878H/+) HSCs, MPP^G/M^ and GMP after 48h culture with 0 or 500ng/ml OSM. Dots show individual mice, bars represent mean ± SEM of *n* = 3.

Next, we examined myeloid differentiation potential. Control and *Dnmt3a*-mutant HSPCs were prospectively isolated from young adult mice and plated into a myeloid methylcellulose media to quantify colony-forming units (CFU). The myeloid methylcellulose media was supplemented with 0, 100 or 500ng/ml recombinant murine OSM. After 7-day culture, we observed no differences in CFU formation across any of the conditions (Figure 2C). This result suggests that control and *Dnmt3a*-mutant HSPCs do not respond to OSM with respect to altering their myeloid differentiation potential. To examine CFU re-plating potential, we continued to passage control and *Dnmt3a*-mutant HSPCs as well as HSCs with 0 or 500ng/ml recombinant murine OSM. As expected, we observed increased CFU re-plating potential from *Dnmt3a*-mutant vs. control HSPCs and HSCs (Figure 2D). However, there was no observed effect of OSM on either control or *Dnmt3a*-mutant CFU re-plating capacity.

A limitation of the above CFU studies is that the effect of OSM is being evaluated in the presence of a full complement of cytokines that drive robust myelo/erythroid cell differentiation (SCF, IL-3, IL-6, and EPO). To examine the effects of OSM on HSPC proliferation in conditions replicating stress, we added 0 or 500ng/ml recombinant murine OSM to previously defined ‘cytokine-poor’ (SCF, G-CSF) and ‘cytokine-rich’ (SCF, GM-CSF, IL-3, IL-11, Flt3L, TPO, EPO) medias^32^ to replicate stress and non-stress conditions, respectively. Control and *Dnmt3a*-mutant HSCs, as well as two progenitor populations (granulocyte/macrophage-primed multipotent progenitors or MPP^G/M^ and granulocyte-macrophage progenitors or GMPs), were prospectively isolated from young adult mice and cultured in cytokine-poor and cytokine-rich medias for 48 hours. After this culture period, total viable cell counts were obtained using flow cytometry. We observed no significant differences in total viable cell counts between genotype and treatment groups for any of the input cell populations (Figure 2E). Together, these data suggest that young control and young *Dnmt3a*-mutant HSPCs do not respond to acute OSM stimulation by altered cell cycling, apoptosis, myeloid differentiation, or proliferation.

### Acute OSM Stimulation Does Not Alter Engraftment Potential or Lineage Output from Young *Dnmt3a*-Mutant HSPCs

As *in vitro* assays do not fully reflect the *in vivo* functional potential of HSPCs, we next tested the hypothesis that recombinant OSM as a single stimulus would be sufficient to promote the selective advantage of young *Dnmt3a*-mutant (R878H/+) HSPCs and HSCs *in vivo*. First, we incubated 50 prospectively isolated HSCs from young adult donor CD45.2+ control or *Dnmt3a*-mutant mice into culture conditions designed to promote HSC self-renewal^33^ supplemented with 500ng/ml recombinant murine OSM or vehicle control (Figure 3A). After 7d culture, the resulting cells in each well were transplanted into lethally irradiated CD45.1+ recipient mice to assess hematopoietic engraftment and lineage potential. At 16wks post-transplant, we observed no significant differences in donor engraftment (% CD45.2+) in the peripheral blood or bone marrow of recipient mice (Figure 3B). Analysis of lineage composition of the peripheral blood graft revealed an increased proportion of B cells from *Dnmt3a*-mutant vs. control vehicle-treated HSCs, as we have previously observed^18^ (Figure 3C). However, we observed no significant differences in peripheral blood lineage composition in recipient mice based on acute OSM stimulation. We also observed no changes in donor-derived bone marrow HSPCs subsets due to acute OSM stimulation, including HSCs, MPP^G/M^, common myeloid progenitors (CMP) and GMP (Figure 3D).

**Figure 3.**
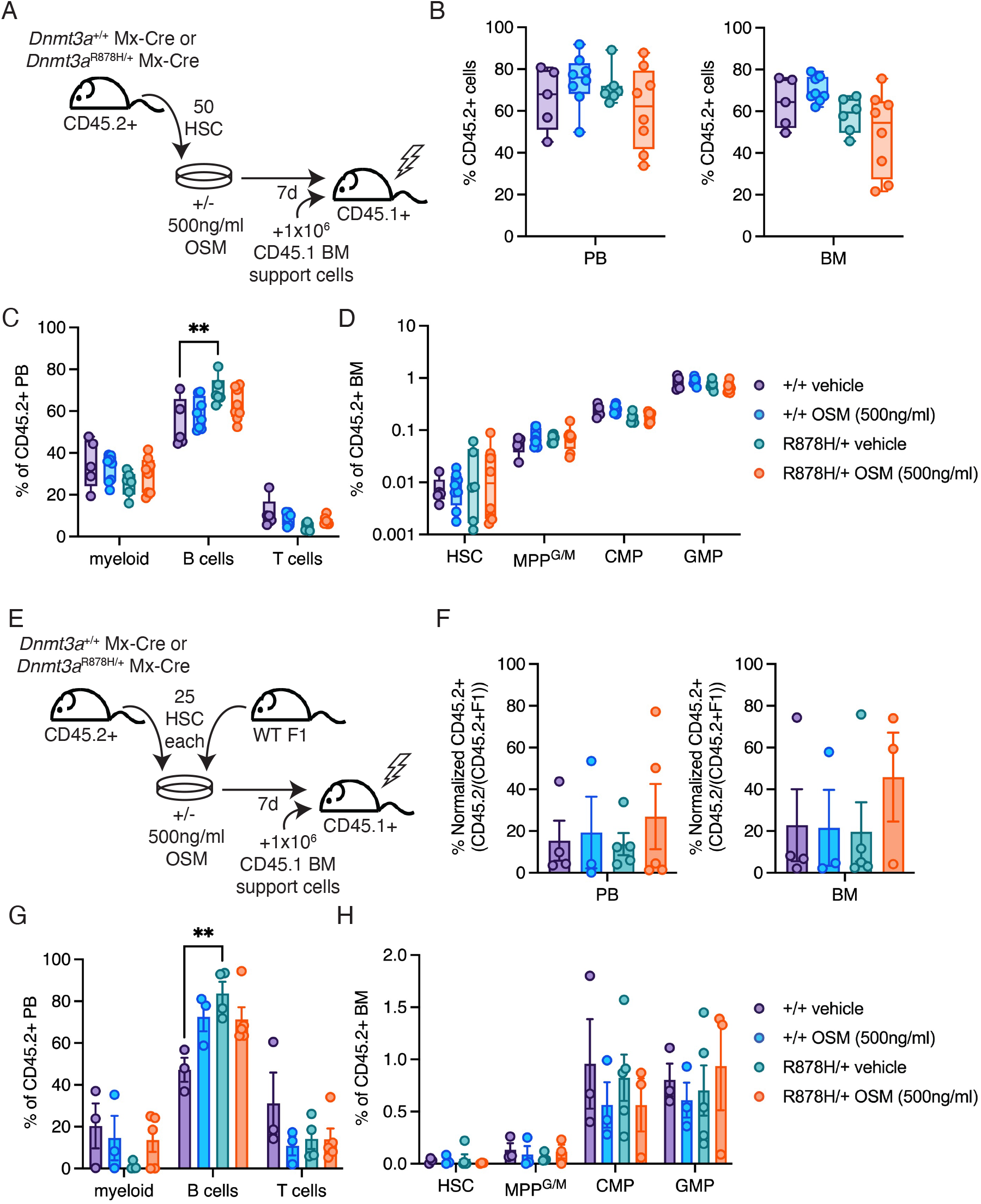
Engraftment potential and lineage output from young *Dnmt3a*-mutant HSPCs is unaltered by acute OSM stimulation. (**A**) Schematic of transplant design. **(B)** Frequency of donor CD45.2^+^ cells in the peripheral blood (PB) and bone marrow (BM) at 16wks post-transplant of control (+/+) and *Dnmt3a*-mutant (R878H/+) HSCs treated with 0 or 500ng/ml OSM for 7d. (**C**) Donor-derived PB lineage (myeloid, B and T cell) frequencies. (**D**) Donor-derived BM hematopoietic stem and progenitor cell frequencies. (**E**) Schematic of competitive transplant design. (**F**) Normalized frequency of donor CD45.2^+^ cells in PB and BM at 16wks post-transplant of control (+/+) and *Dnmt3a*-mutant (R878H/+) HSCs treated with 0 or 500ng/ml OSM for 7d. (**G**) Donor-derived PB lineage (myeloid, B and T cell) frequencies. (**H**) Donor-derived BM hematopoietic stem and progenitor cell frequencies. In all figures, dots show individual mice and bars represent mean ± SEM of *n* = 3-8. \*\**p* < 0.01 by two-way ANOVA with Tukey’s multiple comparisons test.

Next, we utilized a competitive transplantation experimental design to evaluate the effect of OSM on cellular competition between *Dnmt3a*-mutant and control hematopoiesis. We incubated 25 prospectively isolated HSCs from young adult donor CD45.2+ control or *Dnmt3a*-mutant mice with 25 prospectively isolated HSCs from competitor CD45.2+ CD45.1+ (F1 hybrid) mice. These were cultured in the same conditions as above, supplemented with 500ng/ml recombinant murine OSM or vehicle control (Figure 3E). After 7-day culture, the resulting cells in each well were transplanted into lethally irradiated CD45.1+ recipient mice to assess hematopoietic engraftment and lineage potential. At 16wks post-transplant, we observed no significant differences in donor engraftment (% CD45.2+) in the peripheral blood or bone marrow of recipient mice (Figure 3F). Analysis of lineage composition of the peripheral blood graft revealed an increased proportion of B cells from *Dnmt3a*-mutant vs. control vehicle-treated HSCs (Figure 3G), consistent with the findings above. However, we observed no significant differences in peripheral blood lineage composition in recipient mice based on acute OSM stimulation. We also observed no changes in donor-derived bone marrow HSPCs subsets due to acute OSM stimulation, including HSCs, MPP^G/M^, CMP and GMP (Figure 3H). Together, these data suggest that acute OSM stimulation does not lead to changes in engraftment or lineage potential of young *Dnmt3a*-mutant HSPCs in non-competitive or competitive transplant.

### Young *Dnmt3a*-Mutant HSCs are Responsive to Acute OSM Stimulation via STAT3 Phosphorylation and Transcriptional Alterations

Due to the lack of phenotypic alterations associated with recombinant OSM stimulation of young control and young *Dnmt3a*-mutant HSCs *in vitro* and *in vivo*, we evaluated the extent to which these cells have the capacity to directly bind and respond to recombinant murine OSM. We first considered antibody-based assessment of levels of the OSM receptor subunit OSMR on the cell surface. Due to a lack of specific and commercially available anti-mouse OSMR antibodies^34^, we fluorescently labeled recombinant OSM (OSM-AF488) and tested binding and labelling of *Dnmt3a*-mutant and control HSCs, in comparison to negative (no OSM-AF488) and positive (liver cells with OSM-AF488) controls. We observed binding of OSM-AF488 to both control and R878H/+ HSCs (Supplemental Figure 1A), supporting that control and *Dnmt3a*-mutant HSCs are capable of directly binding recombinant murine OSM.

We experimentally evaluated activation of STAT3 by recombinant OSM in *Dnmt3a*-mutant HSPCs, which was predicted by our original data (Figure 1D). We prospectively isolated control and *Dnmt3a*-mutant HSPCs from young adult mice, stimulated *ex vivo* with 500ng/mL of recombinant murine OSM over a time course and evaluated phosphorylation of STAT3 and STAT5 by flow cytometry (Figure 4A). We observed that acute OSM stimulation of *Dnmt3a*-mutant HSPCs resulted in greater pSTAT3 compared to vehicle-stimulated *Dnmt3a*-mutant HSPCs as well as compared to OSM-stimulated control HSPCs after 60min (Figure 4B). No differences were observed in pSTAT3 in any condition after 20min stimulation or 80min stimulation, indicating tight regulation of the OSM-STAT3 signaling response. No differences were observed in pSTAT5 in any condition at any of the tested time points (Supplemental Figure 1B), demonstrating selectivity of acute OSM signaling response towards STAT3 activation in young *Dnmt3a*-mutant HSPCs.

**Figure 4.**
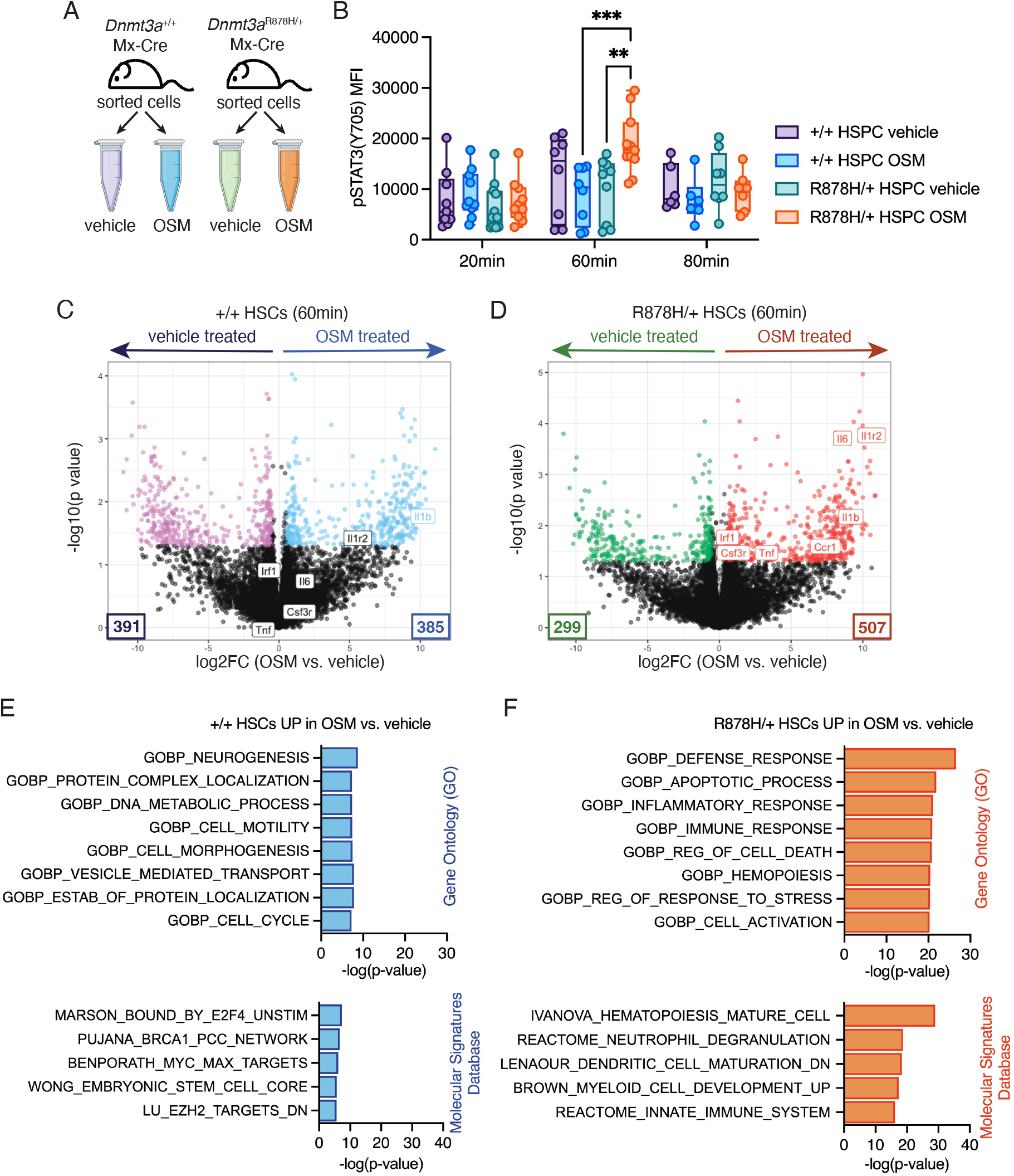
Acute OSM stimulation activates STAT3 phosphorylation and transcriptional responses in young *Dnmt3a*-mutant HSCs. (**A**) Schematic of experimental design. (**B**) Mean fluorescence intensity (MFI) of pSTAT3 (Y705) in control (+/+) and *Dnmt3a*-mutant (R878H/+) HSPCs treated with 0 or 500ng/ml OSM for 20, 60 and 80min. Dots show individual mice and bars represent mean ± SEM of *n* = 6-10. \*\**p* < 0.01; \*\*\**p* < 0.001 by mixed-effects analysis with Tukey’s multiple comparisons test. (**C, D**) Volcano plot showing differential gene expression of (**C**) control (+/+) and (**D**) *Dnmt3a*-mutant (R878H/+) HSCs treated with 0 or 500ng/ml OSM for 60min. *n* = 6 biological replicates. Select genes involved in STAT3 signaling and inflammation are labelled. (**E, F**) Enrichment analysis of significantly differentially expressed genes in (**E**) control (+/+) and (**F**) *Dnmt3a*-mutant (R878H/+) HSCs treated with 500ng/ml vs. 0ng/ml OSM for 60min.

We sought to test the transcriptional consequences of OSM-STAT3 signaling in a more purified control and *Dnmt3a*-mutant HSC population. We prospectively isolated control and *Dnmt3a*-mutant HSCs from young adult mice, stimulated *ex vivo* with 500ng/mL of recombinant murine OSM or vehicle control for 60min and immediately flash froze cell pellets for RNA extraction and RNA-seq. We identified significantly differentially expressed genes comparing OSM- vs. vehicle-treated control HSCs, and OSM- vs. vehicle-treated *Dnmt3a*-mutant HSCs, using *P* < 0.05 and fold change (FC) ± 1.5 cutoffs. This analysis revealed OSM-treated control HSCs had 385 genes increased in expression and 391 genes decreased in expression (Figure 4C). In contrast, OSM-treated *Dnmt3a*-mutant HSCs had 507 genes increased in expression and 299 genes decreased in expression (Figure 4D). Of the 385 OSM-activated genes in control HSCs and the 507 OSM-activated genes in *Dnmt3a*-mutant HSCs, only 19 were overlapping, suggesting a fundamentally distinct transcriptional response of young *Dnmt3a*-mutant HSCs to acute OSM. Delving further into specific gene alterations, we noted that acute OSM-stimulated *Dnmt3a*-mutant HSCs had robust upregulation of several key inflammatory molecules, receptors and response factors including *Il6, Il1b*, *Tnf, Il1r2, Csf3r, Ccr1* and *Irf1*. Apart from *Il1b*, these were not upregulated in OSM-stimulated control HSCs. These data suggests that a 60min stimulation of young *Dnmt3a*-mutant HSCs with recombinant OSM is sufficient to induce transcriptional upregulation of inflammatory cytokines associated with *Dnmt3a*-mutant clonal hematopoiesis.

To further interrogate these transcriptional signatures, we performed enrichment analyses using gene ontology (GO) terms as well as using the molecular signatures database (MSigDB). Acute OSM-stimulated genes in *Dnmt3a*-mutant HSCs were enriched for focused signatures of inflammation, apoptosis, immune defense/stress responses, innate immunity, hematopoiesis, and myeloid cell development (Figure 4F). In contrast, OSM-stimulated genes in control HSCs were enriched for generic signatures of DNA metabolic processes, cell motility and morphogenesis, vesicle-mediated transport, and cell cycle-related processes (Myc/Max targets and E2F4 binding) (Figure 4E). Together, these data support that acute stimulation of *Dnmt3a*-mutant HSCs with recombinant OSM results in OSM binding, STAT3 phosphorylation, and transcriptional activation of an inflammatory gene network including the inflammatory cytokines IL-6, TNFα and IL-1β that are associated with *Dnmt3a*-mutant clonal hematopoiesis in mice and humans.

### *Dnmt3a-*mutant HSCs Upregulate Anti-Inflammatory Genes in Response to Acute OSM Stimulation but not Chronic Stimulation in an Aged Environment Context

We sought to resolve the disconnect we observed between acute OSM-driven STAT3 signaling and transcriptional activation in *Dnmt3a*-mutant (R878H/+) HSCs and the lack of phenotypic changes in *Dnmt3a*-mutant HSCs and HSPCs. Previous studies have found elevated expression of suppressors of inflammation such as *Socs3, Nr4a1* and *Atf3* in CH-mutant HSPCs^35^, and OSM has been reported to stimulate expression of *Socs3* in other cell types^36, 37^. Thus, we hypothesized that acute OSM stimulation of young *Dnmt3a*-mutant HSCs results in enhanced expression of suppressors of inflammation. Interrogating our OSM- vs. vehicle-treated *Dnmt3a*-mutant HSC RNA-seq data revealed that *Socs3, Nr4a1* and *Atf3* were increased in expression after acute OSM stimulation (Figure 5A).

**Figure 5.**
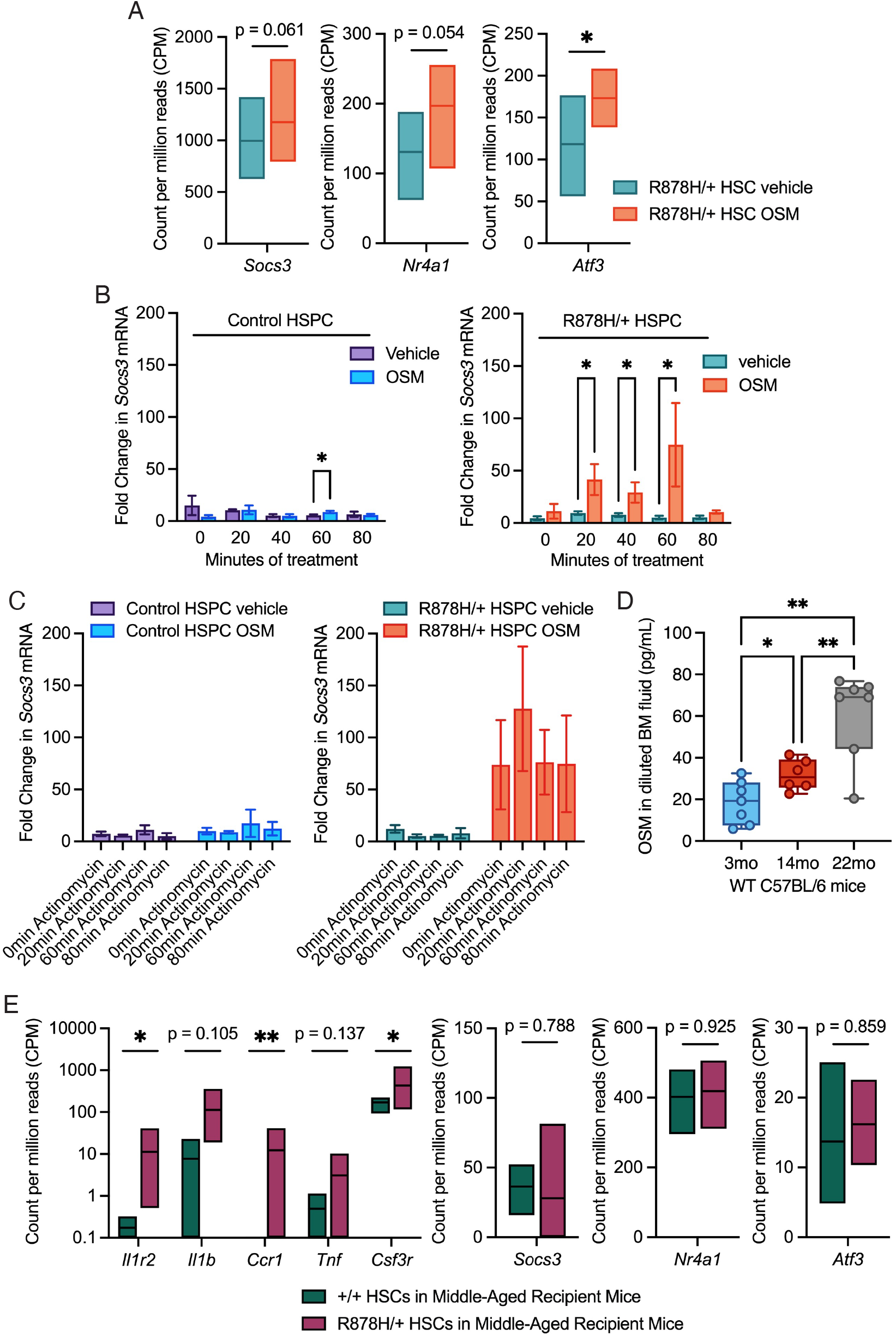
*Dnmt3a*-mutant HSCs exhibit upregulation of anti-inflammatory genes in response to acute OSM stimulation but not in an aging context. (**A**) *Socs3*, *Nr4a1* and *Atf3* expression in control (+/+) and *Dnmt3a*-mutant (R878H/+) HSCs stimulated with 0 or 500ng/ml OSM for 60min. Box plots summarize *n* = 6 replicates per condition. \**p* < 0.05. (**B**) *Socs3* expression in control (+/+) and *Dnmt3a*-mutant (R878H/+) HSPCs stimulated with 0 or 500ng/ml OSM for described amount of time. Bars represent mean ± SEM of *n* = 3-4 replicates. \**p* < 0.05 by multiple-ratio, paired sample t test. (**C**) *Socs3* expression in control (+/+) and *Dnmt3a*-mutant (R878H/+) HSPCs stimulated with 0 or 500ng/ml of OSM for 60min followed by actinomycin D for 0, 20, 60 and 80min. Bars represent mean ± SEM of *n* = 4 replicates. (**D**) OSM concentration in bone marrow fluid in young (3mo), middle-aged (14mo) and old (22mo) wild-type C57BL/6 mice. Dots show individual mice, bars represent mean ± SEM of *n =* 6-7. \**p <* 0.05; \*\**p* < 0.01 by Brown-Forsythe and Welch’s ANOVA with multiple comparisons. (**E**) Expression of inflammatory genes *Il1r2, Il1b, Ccr1, Tnf,* and *Csf3r*, and the anti-inflammatory genes *Socs3, Nr4a1*, and *Atf3* in control (+/+) and *Dnmt3a*-mutant (R878H/+) HSCs after 4mos post-transplant in middle-aged recipient mice. Box plots summarize *n* = 3-4 replicates per condition. \**p* < 0.05; \*\**p* < 0.01 by multiple-ratio, paired sample t test.

To rigorously test the dynamics of transcript induction, we focused on *Socs3*. We prospectively isolated control and R878H/+ HSPCs from young adult mice and stimulated with 0 or 500ng/ml recombinant murine OSM for up to 80min. Cells were flash-frozen for RNA extraction, cDNA synthesis and quantitative real-time PCR for *Socs3*. We observed that OSM stimulation of control HSPCs resulted in a small, but significant, increase in *Socs3* after 60min (Figure 5B). In contrast, OSM stimulation of *Dnmt3a*-mutant HSPCs resulted in robust increase in *Socs3* at 20, 40 and 60min. This result demonstrates that *Dnmt3a*-mutant HSPCs rapidly respond to OSM stimulation by upregulating *Socs3*, at both greater levels and a faster rate compared to control HSPCs.

We also evaluated stability of the *Socs3* mRNA transcript. We prospectively isolated control and *Dnmt3a*-mutant HSPCs from young adult mice and stimulated with 0 or 500ng/ml recombinant murine OSM for 60min, a timepoint we have consistently shown to result in STAT3 phosphorylation and transcriptional responses in *Dnmt3a*-mutant HSPCs. We followed this stimulation by incubating cells for 0, 20, 60 and 80 minutes with Actinomycin D, to pause transcription and allow the study of transcript stability over time^38^. Cells were flash-frozen for RNA extraction, cDNA synthesis and real-time PCR for *Socs3*. Consistent with the above results, *Socs3* was robustly increased in *Dnmt3a*-mutant HSPCs by 60min of OSM stimulation (Figure 5C). Furthermore, we observed that OSM-stimulated *Dnmt3a*-mutant HSPCs maintained increased levels of *Socs3* for up to 80 minutes of Actinomycin D treatment. These results suggest that OSM-stimulated *Dnmt3a*-mutant HSPCs have a robust increase in *Socs3* transcript expression, and that this transcript is stably maintained for up to 140 minutes following exposure to OSM. We posit that *Socs3* activation may suppress STAT3 signaling, silencing transcriptional responses to OSM before functional outcomes are realized.

At the beginning of our study, we discovered a transcriptional program of inflammation response in *Dnmt3a*-mutant compared to control HSCs in middle-aged, but not young, transplant recipient mice (Figure 1B). Thus, we hypothesized that middle-aged mice have elevated ‘chronic’ levels of OSM and that *Dnmt3a*-mutant HSCs in a middle-aged microenvironment lose the capacity to upregulate suppressors of inflammation such as *Socs3, Nr4a1,* and *Atf3*. We collected bone marrow fluid from young adult (3mo), middle-aged (14mo) and old (22mo) wild-type C57BL/6 mice and quantified OSM abundance using ELISA. We observed a significant and progressive increase in the quantity of OSM in the bone marrow fluid from young to middle-aged to old mice (Figure 5D). We examined our published RNA-seq data of control and *Dnmt3a*-mutant HSCs re-isolated from transplanted middle-aged wild-type recipient mice, focusing on the key inflammatory molecules identified in our acute OSM-stimulated *Dnmt3a*-mutant HSCs (Figure 4D) as well as *Socs3, Nr4a1* and *Atf3*. We found that *Dnmt3a*-mutant HSCs compared to control HSCs in transplanted middle-aged recipient mice had robust upregulation of *Il1r2, Il1b*, *Ccr1, Tnf,* and *Csf3r* (Figure 5E). Unlike acute OSM-stimulated *Dnmt3a*-mutant HSCs, we did not observe increased expression of *Socs3, Nr4a1* or *Atf3*.

## DISCUSSION

We have uncovered a role for Oncostatin M (OSM) as a master regulator of an inflammatory cytokine network active in *Dnmt3a*-mutant CH. Our initial discovery was based on transcriptional signatures indicating active OSM signaling in *Dnmt3a*-mutant HSCs specifically in the context of a middle-aged bone marrow microenvironment. In functional experiments, we found that OSM stimulation of young *Dnmt3a*-mutant HSCs did not impact hematopoietic cell function or output *in vitro* or *in vivo*, however, it did result in STAT3 phosphorylation and a transcriptional inflammatory response including upregulation of *Il6, Il1b* and *Tnf*. Focused studies of transcript production and stability revealed a negative feedback mechanism in young *Dnmt3a*-mutant HSCs where OSM signaling results in increased transcription and transcript stability of anti-inflammatory genes including *Socs3*, *Nr4a1* and *Atf3* (Supplemental Figure 2). In the context of a middle-aged bone marrow microenvironment, which has chronically increased levels of OSM, we find that *Dnmt3a*-mutant HSCs upregulate an inflammatory response transcriptional signature which is not accompanied by upregulation of anti-inflammatory genes. Together, our work suggests that OSM is upstream of an inflammatory cytokine network in *Dnmt3a*-mutant HSCs. We speculate that chronic inflammation or exposure to ligands such as OSM with aging exhausts the regulatory mechanisms present in young *Dnmt3a*-mutant HSCs that resolve inflammatory states.

OSM is an IL-6 family cytokine known to be involved in the immunopathogenesis of colon cancer, breast cancer, pancreatic cancer, myeloma and hepatoblastoma^34^. Whereas IL-6 represents one of the most studied cytokines to date, the physiological activities of OSM are less well known. OSM is predominantly produced by T lymphocytes, macrophages, and neutrophils^19^, and in mice, signals through the heterodimeric receptor gp130/OSMR^39^. OSM is a strong inducer of JAK/STAT signaling leading to activation of STAT3 and STAT5^40, 41^. OSM is an important regulator of the bone marrow microenvironment in both steady state and in regeneration after injury^42, 43^ with endothelial and mesenchymal cells being major cell types expressing OSMR^34^, and plays a role in HSC mobilization via its effects on non-hematopoietic cells in the bone marrow microenvironment^34^. These data and models support a role for OSM as a cytokine produced by mature hematopoietic cells that impacts functionality of non-hematopoietic cells in the bone marrow microenvironment. Previous studies performed in young adult mice have shown that *Osmr* is not detected in HSPCs or myeloid cells^34^ and thus it has been suggested that HSCs and their progeny do not directly respond to OSM. However, *Osmr* is one of the most upregulated transcripts in old HSCs compared to young HSCs across six independent datasets^44^ suggesting that in certain contexts, such as aging, HSCs and their progeny gain the capacity to respond to OSM. Our data show that HSCs, in the context of a CH-relevant mutation in *Dnmt3a*, do have the capacity to phosphorylate STAT3 and undergo transcriptional alterations in response to OSM.

The consequences of OSM signaling are also context dependent, resulting in pro-inflammatory as well as anti-inflammatory/anti-proliferative outcomes^40^. This is consistent with our data in young *Dnmt3a*-mutant HSCs where we show that both pro-inflammatory cytokines as well as anti-inflammatory molecules are expressed in response to OSM. This anti-inflammatory negative feedback loop may be a conserved mechanism facilitating the selective advantage of mutant HSCs in CH. Recent work using a zebrafish model of human *ASXL1*-mutant CH found a protective response to pro-inflammatory cytokines in HSPCs via upregulation of *socs3a, atf3* and *nr4a1*^35^. Without the capacity to upregulate *nr4a1* and *nr4a3*, the *Asxl1*-mutant HSPCs lost their self-renewal capacity and selective growth advantage. In addition to induction of these anti-inflammatory transcripts that work in part through silencing STAT3 activation, complementary mechanisms may allow mutant HSCs to survive in a chronic inflammatory environment associated with aging. For example, in *Tet2*-mutant CH, HSPCs with hyperactivated SHP2-STAT3 signaling downregulate the apoptotic protein Bim via the anti-apoptotic long non-coding RNA *Morrbid*^37^. We find that in the context of a middle-aged BM microenvironment, which has chronically elevated levels of OSM, *Dnmt3a*-mutant HSCs no longer upregulate key anti-inflammatory genes. We speculate that chronic inflammation prompts epigenetic rewiring or selection of *Dnmt3a*-mutant HSCs such that they lose the ability to silence these signals, which merits further experimentation and exploration.

Identification of OSM as a master regulator of inflammatory cytokine production opens the possibly of therapeutic interventions targeting OSM to reduce inflammation in aging-associated *Dnmt3a*-mutant CH. To provide evidence for blocking OSM:OSMRβ signaling in a mouse model of sepsis was found to reduce serum levels of IL-6 and TNFα and prolong mouse survival^45^ and human monoclonal OSMRβ-targeting antibodies such as Vixarelimab are utilized clinically to manage such conditions as prurigo nodularis, a debilitating chronic skin disease^46^. Alternatively, identifying a transcriptional feedback loop that silences pro-inflammatory responses in *Dnmt3a*-mutant HSCs, but is dysfunctional in the context of aging, also represents a targetable mechanism for therapeutic intervention. Re-engagement of this negative feedback loop by transient activation of *Socs3* or other anti-inflammatory factors may help to suppress inflammatory myeloid cell production that is a major contributor to CH-associated diseases including cardiovascular disease and myeloid malignancy.

## AUTHOR CONTRIBUTIONS

L.S.S. and J.J.T. conceptualized the project and designed experiments. L.S.S. performed experiments and analyzed data, with assistance from K.A.Y., N.B. and K.D.M. RNA-seq data was analyzed and graphed by L.S.S. and T.M.S. The manuscript was written by L.S.S. and J.J.T. All authors edited the manuscript.

## ACKNOWLEDGEMENTS

This work was supported by National Institutes of Health grants R01DK118072, R01AG069010, U01AG077925, and an EvansMDS Discovery Research Grant to J.J.T. This work was supported in part by the NIH/NCI Cancer Center Support Grant P30CA034196. J.J.T. was supported by a Leukemia & Lymphoma Society Scholar Award and The Dattels Family Endowed Chair. L.S.S. was supported by F31DK127573 and The Tufts University Scheer-Tomasso Fund philanthropic gift. We thank all members of the Trowbridge Lab for experimental support and manuscript editing. We thank the Scientific Services at The Jackson Laboratory including flow cytometry and genome technologies. We thank Drs. Carol Bult, Ryan Tewhey, Cliff Rosen, and Phil Hinds for their critical input into this work.

## MATERIALS AND METHODS

### Animals

C57BL/6J (JAX:000664) and B6.SJL-*PtprcaPepcb*/BoyJ^47^ (JAX:002014, referred to as CD45.1^+^) mice were obtained from, and aged within, The Jackson Laboratory. *Dnmt3a*^fl-R878H/+^ mice (JAX:032289) were crossed to B6.Cg-Tg(Mx1-cre)1Cgn/J mice (JAX:003556, referred to as Mx-Cre). In all experiments, control (+/+) mice carried a single copy of Mx-Cre allele. The Jackson Laboratory’s Institutional Animal Care and Use Committee approved all experiments. To induce Mx-Cre, mice were intraperitoneally injected once every other day for five total injections with 15 mg/kg high molecular weight polyinosinic-polycytidylic acid (polyI:C) (InvivoGen). In all experiments, mice were used >4 weeks following polyI:C administration.

### Peripheral Blood Analysis

Blood was collected from mice via the retro-orbital sinus and red blood cells were lysed prior to staining with the following fluorochrome-conjugated antibodies: BV650 CD45.1 (BioLegend clone A20), AlexaFluor700 CD45.2 (BioLegend clone 104), BUV496 B220 (BD Biosciences clone RA3-6B2), PerCP-Cy5.5 CD3e (BioLegend clone 145-2C11), APC-Cy7 CD11b (BioLegend clone M1/70), APC Ly6g (BioLegend clone 1A8), BV605 Ly6c (BioLegend clone HK1.4), BV421 Ter-119 (BioLegend clone TER-119), PE-Cy7 F4/80 (Invitrogen BM8). Data was collected using a LSRII (BD Biosciences) and analyzed using FlowJo V10 (BD Biosciences).

### Isolation and Phenotyping of Hematopoietic Stem and Progenitor Cells

BM cells were isolated from pooled and crushed femurs, tibiae, iliac crests, sternums, forepaws, and spinal columns of individual mice. BM mononuclear cells (MNCs) were isolated by Ficoll-Paque (GE Healthcare Life Sciences) density centrifugation or 1X RBC Lysis Buffer (eBioscience) and stained with a combination of fluorochrome-conjugated antibodies: c-Kit (BD Biosciences, BioLegend clone 2B8), Sca-1 (BioLegend clone D7), CD150 (BioLegend clone TC15-12F12.2), CD48 (BioLegend clone HM48-1), CD34 (BD Biosciences clone RAM34), FLT3 (BioLegend clone A2F10), CD11b (BioLegend clone M1/70), mature lineage (Lin) marker mix (B220 (BD Biosciences, BioLegend clone RA3-6B2), CD4 (BioLegend clone RM4-5), CD5 (BioLegend clone 53-7.3), CD8a (Biosciences, BioLegend clone 53-6.7), Ter-119 (BioLegend clone TER-119), and Gr-1 (BioLegend, Invitrogen clone RB6-8C5)), and the viability stain propidium iodide (PI) or 4′,6-diamidino-2-phenylindole (DAPI). For transplants CD45.1 (BioLegend clone A20), and CD45.2 (BioLegend clone 104) antibodies were used to distinguish genotypes of donor and recipient mice. The following cell surface markers were used to isolate or phenotype HSCs: Lin-Sca-1+ c-Kit+ Flt3-CD150+ CD48-, HSPCs: Lin-Sca-1+ c-Kit+, MPP^G/M^: Lin-Sca-1+ c-Kit+ Flt3-CD150-CD48+, CMP: Lin-Sca-1-c-Kit+ CD34+ FcγR-, GMP: Lin-Sca-1-c-Kit+ CD34+ FcγR+. Data was collected using a BD FACSymphony A5 or cells were prospectively isolated using a FACSymphony S6 (BD Biosciences). All flow cytometry data was analyzed using FlowJo V10.

### Cell Cycle Analysis

5,000 HSPCs were sorted directly into TC-treated 96-well plates (Falcon) containing StemSpan™ SFEM II (Stemcell Technologies) with Pen-Strep (Fisher Scientific) and SCF (100 ng/ml, BioLegend), TPO (50 ng/μl, Peprotech), with or without OSM (500 ng/ml, BioLegend) for 24hrs at 37°C and 5% CO2. Cells were stained with Ghost UV450 viability dye (Cytek Biosciences) and fixed with the FIX & PERM™ Cell Permeabilization Kit (Invitrogen) following the manufacturer’s protocol. Following fixation cells were stained with FITC anti-mouse/human Ki-67 (BioLegend) and DAPI. Data was collected on a BD FACSymphony A5.

### Apoptosis Analysis

5,000 HSPCs were sorted directly into 96-well plates containing SFEMII with Pen-Strep and SCF (100 ng/ml), TPO (50 ng/μl), with or without OSM (500 ng/ml) for 24hrs at 37°C and 5% CO2. Cells were stained with Annexin V and Propidium iodide using the Annexin A5 Apoptosis Detection Kit (BioLegend). Data was collected on a FACSymphony A5 (BD Biosciences).

### Colony-Forming Unit (CFU) Assay

HSCs or HSPCs were isolated and plated in MethoCult GF M3434 (StemCell Technologies), with or without OSM (500 ng/ml) and cultured at 37°C and 5% CO2. Colonies were scored between 6- and 14-days post-plating using a Nikon Eclipse TS100 inverted microscope. For serial replating, cells were harvested by washing the plates and 10,000 cells (from HSCs) or 15,000 cells (from HSPCs) were replated into fresh MethoCult GF M3434 with or without OSM (500 ng/ml).

### In Vitro Culture with Cytokine-Rich and -Poor Media

We followed a published protocol to generate cytokine-rich and cytokine-poor medias^32^. Briefly, 500 cells were sorted directly into 96-well plates in 200μl of IMDM containing 5% FBS, 50 U/ml penicillin, 50μg/ml streptomycin, 2mM L-glutamine, 0.1mM non-essential amino acids, 1mM sodium pyruvate and 50μM 2-mercaptoethanol. For cytokine-rich media, this was supplemented with SCF (25 ng/ml), TPO (25 ng/ml), Flt3L (25 ng/ml), IL-11 (25 ng/ml), IL-3 (10 ng/ml), GM-CSF (10 ng/ml) and EPO (4 U/ml) (all from PeproTech). For cytokine-poor media, this was supplemented with SCF (25 ng/ml) and G-CSF (25 ng/ml, PeproTech). PBS or OSM was added at 500ng/ml to both medias. After 48hr culture at 37°C and 5% CO2, cells were harvested, DAPI was used to determine live cells and data collected on a FACSymphony A5.

### PVA Culture and In Vivo Transplantation

For the non-competitive PVA culture and transplant experiment, 50 CD45.2^+^ HSCs were sorted into a 96-well plate with Ham’s F12 media containing 1X pen/strep/glutamine (Gibco), 10 mmol/l HEPES (Gibco), 1X insulin/transferrin/selenium/ethanolamine (Gibco), 100 ng/ml rmTPO (BioLegend), 10 ng/ml rmSCF (STEMCELL Technologies) and 1 mg/ml polyvinyl alcohol (Sigma), with or without 500 ng/ml rmOSM, and cultured for 7 days at 37°C and 5% CO2, as previously described^33^. 500 ng/ml OSM or vehicle was added to the cultures on days 4 and 6. On day 7, the wells were harvested, mixed with 10^6^ CD45.1^+^ BM MNCs, and transplanted into young, lethally irradiated (12Gy gamma irradiation, split dose) CD45.1^+^ recipients. PB and BM data were collected at 6 months after transplant using a BD LSR II instrument.

For competitive PVA culture and transplant, 25 CD45.2^+^ HSCs from control or R878H/+ donors (CD45.2^+^) and 25 CD45.1^+^CD45.2^+^ HSCs from WT F1 mice were sorted into a 96-well plate with Ham’s F12 media containing 1X pen/strep/glutamine (Gibco), 10 mmol/l HEPES (Gibco), 1X insulin/transferrin/selenium/ethanolamine (Gibco), 100 ng/ml rmTPO (BioLegend), 10ng/ml rmSCF (STEMCELL Technologies) and 1mg/ml polyvinyl alcohol (Sigma), with or without 500 ng/ml rmOSM, and cultured for 7 days at 37°C and 5% CO2, as previously described^33^. 500 ng/ml rmOSM or vehicle was added to the cultures on days 4 and 6. On day 7, the wells were harvested, mixed with 10^6^ CD45.1^+^ BM MNCs, and transplanted into young, lethally irradiated (12 Gy gamma irradiation, split dose) CD45.1^+^ recipients. PB and BM were collected at 6 months after transplant using a BD LSR II instrument.

### Fluorescent OSM Binding

100 μg rmOSM (BioLegend) was concentrated by centrifugation (Amicon Ultra-0.5 Centrifugal Filter Unit) and fluorescently labeled using the AlexaFluor™ 488 Antibody Labeling Kit following the manufacturers protocol. 10^7^ WBM cells were treated with FC block then stained for 30min with PBS or fluorescently labelled rmOSM at 37°C for 30min. After 30min, cells were stained with an antibody cocktail for identification of HSCs and HSPCs as detailed above. Cells were washed and ran on a BD FACSymphony A5 SE. As positive control, a single cell suspension of liver cells was prepared using the Miltenyi Liver Kit (MiltenyiBiotec).

### Phospho-Flow Cytometry

1000 LSK cells were sorted into StemSpan SFEM II media (STEMCELL Technologies). Cells were pelleted and resuspended in StemSpan SFEM II with or without 500 ng/ml rmOSM and incubated at 37°C for 20, 60 or 80min. Cells were then fixed using 16% PFA for 10min at room temperature followed by ice-cold acetone for 10min. Cells were then pelleted, washed and stained with AlexaFluor 488 Mouse Anti-Stat3 (Tyr705) (D3A7) XP Rabbit mAb (Cell Signaling Technologies) or PE-Cy7 Mouse Anti-Stat5 (pY694) (BD) for 30min at room temperature before data collection using a BD FACSymphony A5 SE.

### RNA Sequencing

2,000 HSCs were sorted into StemSpan SFEM II media with TPO (50 ng/ml) and SCF (100 ng/ml). PBS or OSM (500 ng/ml) was added to each well and incubated at 37°C for 60min. Total RNA was isolated from flash-frozen pellets using the RNeasy Micro kit (Qiagen) including the optional DNase digest step. RNA concentration and quality were assessed using the RNA 6000 Pico Assay (Agilent Technologies). Libraries were constructed using the SMARTer Stranded Total RNA-Seq Kit v2-Pico (Takara), according to the manufacturer’s protocol. Library concentration and quality were assessed using the D5000 ScreenTape (Agilent Technologies) and Qubit dsDNA HS Assay (ThermoFisher). Libraries were sequenced (2021) 75 bp paired-end on an Illumina NextSeq 500 using the High Output Reagent Kit v2.5 (2022)150 bp paired-end on an Illumina NovaSeq 6000 using the S4 Reagent Kit v1.5 both at a sequencing depth of >35 million reads per sample. Trimmed alignment files were processed using RSEM (v1.3.3). Alignment was completed using Bowtie 2 (v2.4.1). Expected read counts per gene produced by RSEM were rounded to integer values, filtered to include only genes that had at least two samples within a sample group having a counts per million reads >1, and passed to R (v4.1.3) and edgeR (v3.36.0) for differential expression analysis. A negative binomial generalized log-linear model was fit to the read counts for each gene. The dispersion trend was estimated by Cox-Reid approximate profile likelihood followed by empirical Bayes estimate of the negative binomial dispersion parameter for each tag, with expression levels specified by a log-linear model. Likelihood ratio tests for coefficient contrasts in the linear model were evaluated producing a p-value per contrast. The Benjamini and Hochberg’s algorithm (P value adjustment) was used to control the false discovery rate (FDR). Differentially expressed genes were investigated for overlap with published datasets using Gene Set Enrichment Analysis, and upstream regulators were predicted using Ingenuity Pathway Analysis software. Features with fold change (FC) > 1.5 or < -1.5, and *P* < 0.05, were declared significantly differentially expressed.

### Socs3 mRNA Expression Assays

For *Socs3* expression assays, HSPCs were sorted into 750 μl of StemSpan SFEM II with vehicle or 500 ng/ml OSM and incubated at 37°C for 20, 40, 60 or 80min. Following each incubation, actinomycin D (Sigma) was added and cells were incubated for an additional 20min at 37°C. Samples were pelleted and flash frozen. For *Socs3* stability assays, HSPCs were sorted into 750 μl of StemSpan SFEM II with vehicle or 500 ng/ml OSM and incubated at 37°C for 60min. Actinomycin D was added for 20, 40, 60 or 80min at 37°C. Samples were pelleted and flash frozen. From all samples, RNA was isolated using the RNeasy Microkit (Qiagen) and cDNA was made using the qPCR Bio cDNA synthesis kit (PCR Biosystems). Quantitative PCR was performed using Power SYBR on the QuantStudio 7 Real-Time PCR System (ThermoFisher Scientific). mRNA expression levels were calculated relative to the housekeeping gene, *B2m*.

### ELISA

Bone marrow fluid was collected from 3mo, 14mo, and 22mo C57BL/6J mice by needle flushing of femurs with 200 μl PBS. OSM concentration was determined using the Quantikine ELISA for mouse Oncostatin M (R&D Systems) using a Spectramax i3 (Molecular Devices) plate reader.

### Statistical Analysis

All statistical tests including evaluation of the normal distribution of data and examination of variance between groups were performed using Prism 9 software (GraphPad). Supplemental Figure 2 was created using BioRender.com licensed to The Jackson Laboratory.

### Data Availability

Raw RNA-seq data is available at the Gene Expression Omnibus (GEO) under accession number GSE236693.

---

**Supplemental Figure 1.**
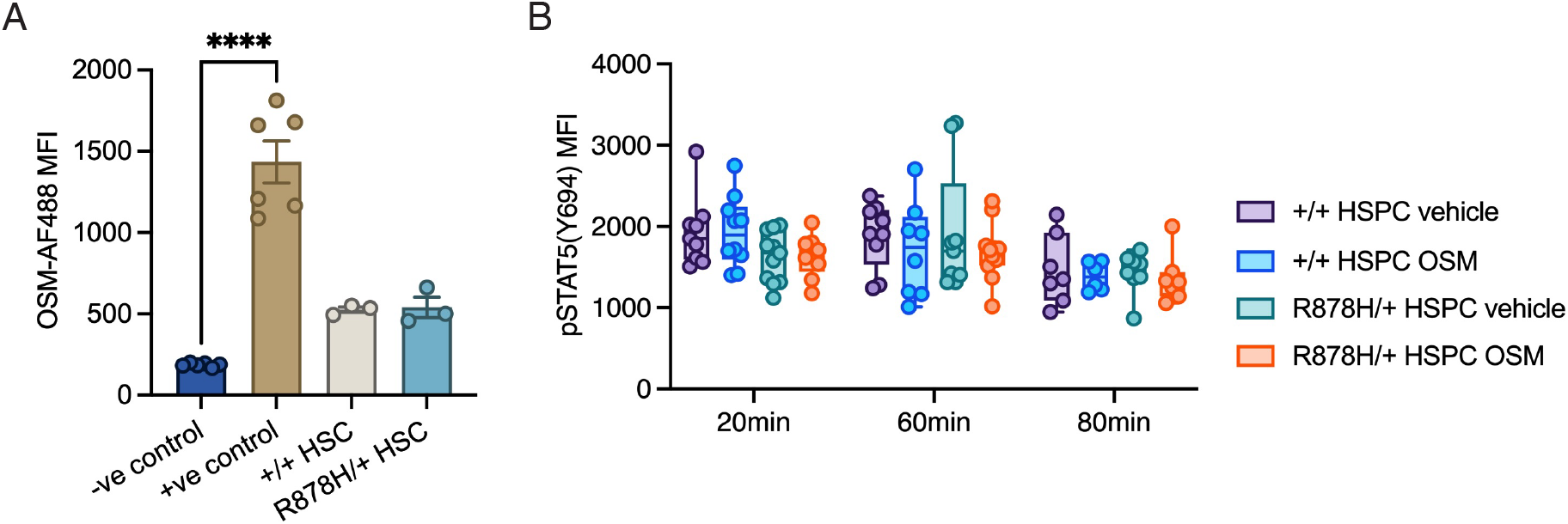
Binding of OSM to control and *Dnmt3a*-mutant HSPCs and induction of pSTAT5. (**A**) Mean fluorescence intensity (MFI) of fluorescently labeled OSM (OSM-AF488) in negative control (no OSM), positive control (liver), control (+/+) and *Dnmt3a*-mutant (R878H/+) HSCs. Dots show individual mice, bars represent mean ± SEM of *n* = 3-6. \*\*\*\**p* < 0.0001 by one-way ANOVA with Dunnett’s T3 multiple comparisons test. (**B**) MFI of pSTAT5 (Y694) in control (+/+) and *Dnmt3a*-mutant (R878H/+) HSPCs treated with 0 or 500ng/ml OSM for 20, 60 and 80min. Dots show individual mice and bars represent mean ± SEM of *n* = 6-10. \*\**p* < 0.01; \*\*\**p* < 0.001 by mixed-effects analysis with Tukey’s multiple comparisons test.

**Supplemental Figure 2.**
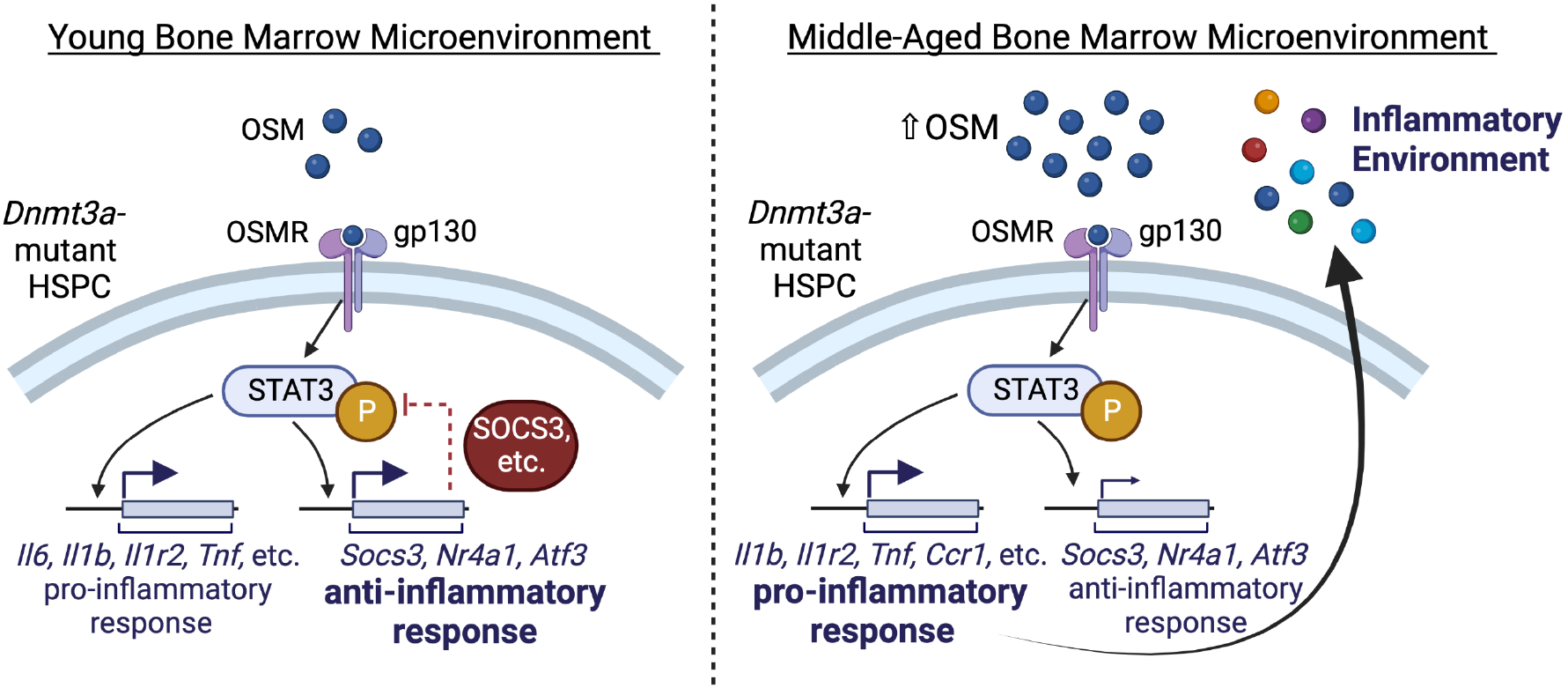
Working model of OSM signaling in *Dnmt3a*-mutant HSPCs in the context of the young and middle-aged BM environments. Created with BioRender.com licensed by The Jackson Laboratory.

